# Childhood Trauma Severity Compounds Reward Anticipation Disruption in Substance Use Disorders

**DOI:** 10.64898/2026.11.13.766203

**Authors:** Samia A. Aguilar, Sarah I. Qundes, Rayus Kuplicki, Robin L. Aupperle, Jennifer L. Stewart, Maëlle C.M. Gueguen

## Abstract

**Background:** Substance use disorders (SUD) are associated with reward circuit disruptions and high rates of trauma. Trauma has been found to reduce propensity to engage frontal, limbic, and striatal regions during reward anticipation. However, the interactive effect of trauma and SUD on anticipatory reward circuitry remains unclear.

**Methods:** To address this critical research gap, treatment-enrolled individuals with at least one SUD (*N* = 150; 78 female) completed questionnaires assessing childhood trauma and current posttraumatic stress disorder (PTSD) symptoms and a monetary incentive delay (MID) task during functional magnetic resonance imaging. The MID task presented participants with cues predicting the delivery of no, small, and large monetary gains and avoidance of equally weighted losses. Linear mixed effects models examined the effect of trauma severity or PTSD on percent signal change for anticipation of monetary gains, extracted from a-priori striatal, prefrontal, and mesolimbic anatomic regions of interest.

**Results:** 70.7% of individuals experienced significant childhood trauma and 63.3% met self-report symptom thresholds for probable PTSD. Current PTSD symptoms were positively associated with emotion dysregulation, emotional impulsivity, and drug craving, but were not related to anticipatory BOLD signal. Higher childhood trauma severity (but not current PTSD severity) was associated with lower anticipatory responses within the nucleus accumbens, caudate, dorsomedial prefrontal cortex, amygdala, anterior cingulate cortex, and anterior insula to both gain and loss cues.

**Conclusions:** Childhood adversity is associated with blunted reward anticipation across striatal, prefrontal, and limbic circuitry. This highlights the importance of considering childhood trauma severity in SUD treatment, which may require targeted interventions beyond standard cognitive behavioral therapy to restore normative salience processing, such as striatal neuromodulation.

## Introduction

Substance use disorders (SUD) are a major public health concern marked by chronic relapse, substantial morbidity, and significant psychosocial burden (1). A key feature of SUD is disruption in reward anticipation, including reduced sensitivity to non-drug rewards and heightened cue-driven motivation for drug-related stimuli (2). During reward anticipation, frontostriatal and insular regions serve distinct but complementary roles: the nucleus accumbens [NAc]) is involved in reward prediction and motivational salience, the caudate and putamen contribute to action-outcome learning and habit formation, the medial prefrontal cortex (PFC) and anterior cingulate cortex (ACC) exert top-down regulatory control over reward-driven behavior, and the insula integrates interoceptive signals that inform affective appraisal of anticipated outcomes.

Meta-analytic work in healthy individuals ((3); (4); (5)) indicates that gain anticipation elicits NAc, caudate, putamen, insula, mPFC, and ACC activation during the monetary incentive delay (MID) task, with the extent of activation scaling parametrically with reward magnitude, producing a characteristic pattern of greater responses to high relative to low incentive conditions and minimal responses to no-incentive trials (6). The MID task has proven particularly valuable for characterizing reward anticipation deficits in SUD, as it reliably dissociates anticipatory from consummatory reward processing across clinical populations (7). Although gain anticipation has been the primary focus of this literature, the MID task also includes loss-avoidance trials, and anticipatory responses to potential loss are increasingly recognized as relevant to both SUD and trauma-related psychopathology, given the role of negative reinforcement and heightened threat sensitivity in each condition. In contrast to this pattern observed in healthy samples, meta-analyses indicate that individuals with SUD experience a blunting of the ventral striatal reward anticipation response (8) that is also evident within meso-cortical circuits (including the insula and mPFC (9)). While both anticipatory and consummatory phases of reward processing are disrupted in SUD, blunting of the anticipatory response is particularly robust and is thought to drive the motivational deficits underlying compulsive drug seeking ((8); (7)), as blunted responsivity to non-drug rewards, combined with heightened reactivity to drug-related cues, may narrow the range of experiences capable of generating motivational salience and reinforce continued reliance on drug use as a primary source of reward.

While interventions targeting striatal, insula, and mPFC upregulation to non-drug rewards may be beneficial for targeting reward impairments in SUD, critical research gaps still exist. First, as MID neuroimaging studies have largely centered on individuals with alcohol, nicotine and cocaine use disorders, it is unknown whether these findings also extend to individuals with methamphetamine and/or opioid use disorders, representing the current dual drug epidemics in the United States today ((10); (11)). Second, childhood trauma exposure and posttraumatic stress disorder (PTSD) symptoms are highly prevalent among individuals with SUD, with one treatment-seeking cohort reporting lifetime trauma exposure in 97% and current PTSD in 37% of patients (12); while there is a growing literature on the relationship between trauma and alterations within reward circuitry, the interaction between SUD and trauma on non-drug reward circuitry has yet to be clarified, particularly for individuals with methamphetamine and/or opioids as their drug(s) of choice. Childhood trauma may carry particular developmental significance in this context, as early-life adversity occurs during sensitive periods of neuroplasticity and has been linked to lasting changes in reward and stress circuitry, yet its contribution to reward anticipation deficits in SUD remains comparatively understudied relative to current PTSD symptomatology.

Childhood maltreatment is specifically associated with amygdala hyperreactivity and altered dorsolateral PFC (dlPFC) and mPFC signaling (28), as well as lasting alterations in corticostriatal and limbic circuits (including posterior cingulate cortex (PCC) involved in motivation and affective valuation ((29); (30); (31); (32); (33)). Early adversity has been associated with blunted striatal responsivity and diminished mesolimbic reward sensitivity in adulthood, neurobiological signatures consistent with a developmentally embedded hyposensitivity to natural reinforcers ((34); (35); (36)).

PTSD is associated with altered (with reports of both blunted and exaggerated activation) responses in the striatum, amygdala, and mPFC during MID gain anticipation ((13); (14)). Although the amygdala is not consistently identified in meta-analyses of healthy or SUD samples during reward anticipation, it is implicated in threat processing and has been shown to play a part in assigning motivational salience to emotionally significant cues, encoding both positive and negative emotional valence and value ((15); (16)). Moreover, the high comorbidity of PTSD and SUD is frequently driven by a negative reinforcement cycle ((17); (18)), in which trauma-related distress motivates drug use as a means of emotional escape or relief. Central to this cycle are difficulties in emotion regulation and the capacity to modulate affective states in adaptive ways, which are robustly elevated in both PTSD and SUD and have been linked to greater severity of trauma symptoms and increased substance use as a coping strategy (18). Compounding this, impulsivity, particularly negative urgency, has been identified as a transdiagnostic vulnerability factor that amplifies maladaptive responding under emotional distress in both trauma-exposed and substance-using populations ((19); (20)). Finally, craving and desire for drug use represent a key motivational component of this cycle, reflecting the degree to which drug cues act as reinforcers in driving behavior ((21); (22); (23)).

Together, these difficulties in emotion regulation, impulsivity, and craving may compound trauma-related alterations on brain regions involved in reward anticipation (24), further aggravated by a dysregulation of prefrontal top-down control, sustained hyperarousal, and disrupted cognitive-affective integration processes ((25); (26); (27)). Taken together, extant findings raise the possibility that childhood trauma history and current PTSD symptoms may exacerbate the blunted striatal and mPFC reward-anticipation signal already observed in SUD, an interactive hypothesis that has not yet been directly tested.

The current study aims to address these gaps by examining neural responses to gain and loss anticipation for individuals with SUD (primary methamphetamine and/or opioid use disorders) who completed the MID task during functional magnetic resonance imaging (fMRI) and questionnaires capturing PTSD symptomatology and trauma history. Our primary aims centered on the impact of (1) childhood trauma on behavioral/clinical measures; and (2) childhood trauma severity on reward anticipation.

## Methods and Materials

### Participants

A total of 150 right-handed participants (78F, 72M) between 18 and 55 years of age were enrolled between January 2019 and November 2022 at the Laureate Institute for Brain Research after providing informed consent (Western IRB #20101611). Exclusion criteria included psychotic disorders, significant neurological illness such as epilepsy or moderate-to-severe traumatic brain injury, and the presence of MRI contraindications including metallic implants, pacemakers, pregnancy, or severe claustrophobia. Participants experiencing severe drug withdrawal or acute substance intoxication were also excluded. Recruitment occurred through treatment centers, community advertisements, and clinician referrals, and written informed consent was obtained prior to any study procedures. Key information on demographics and comorbid psychiatric diagnoses is provided in **Table 1**. For a full description of the sample, see **Table S1**.

**Table 1:**
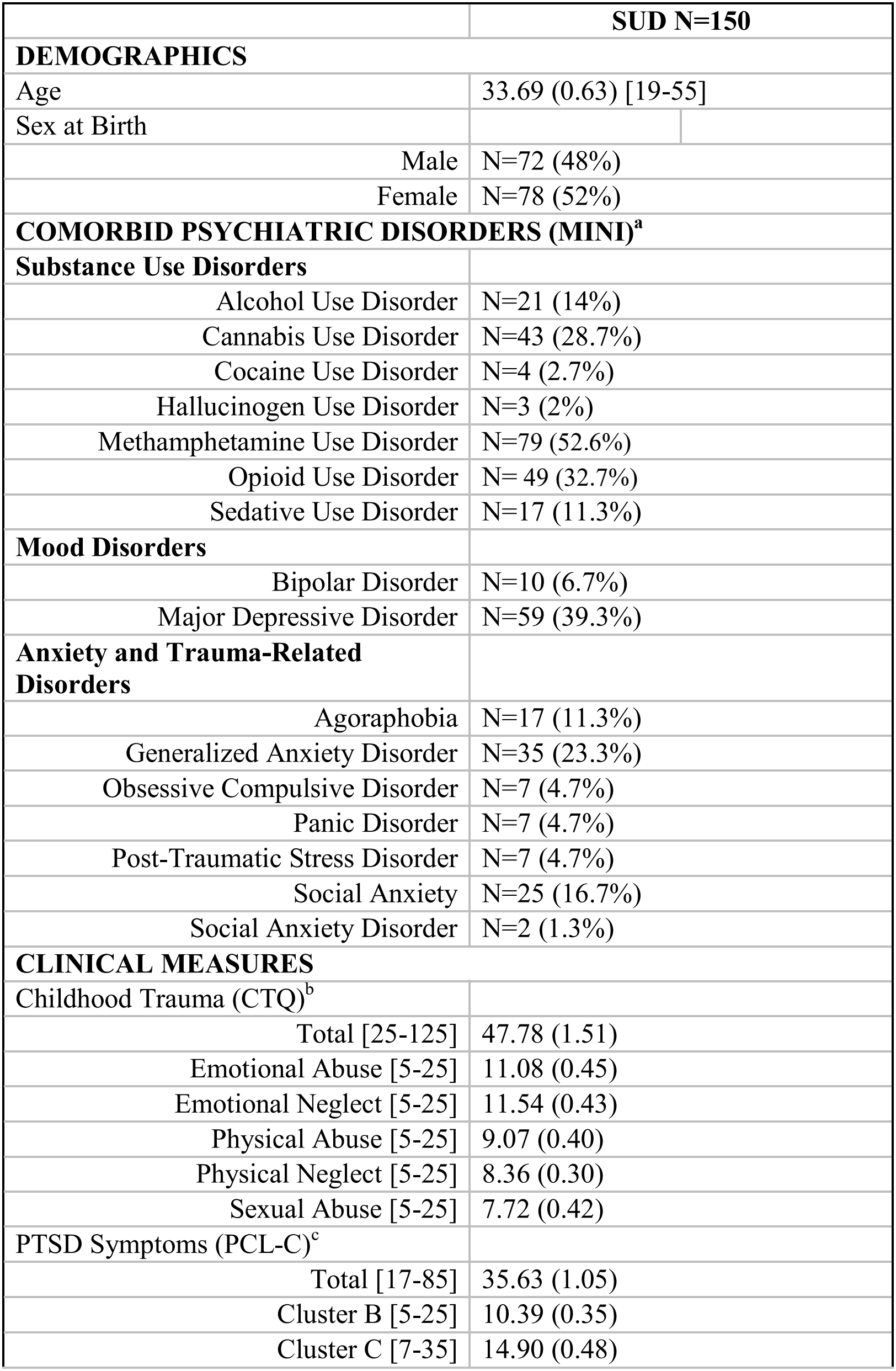
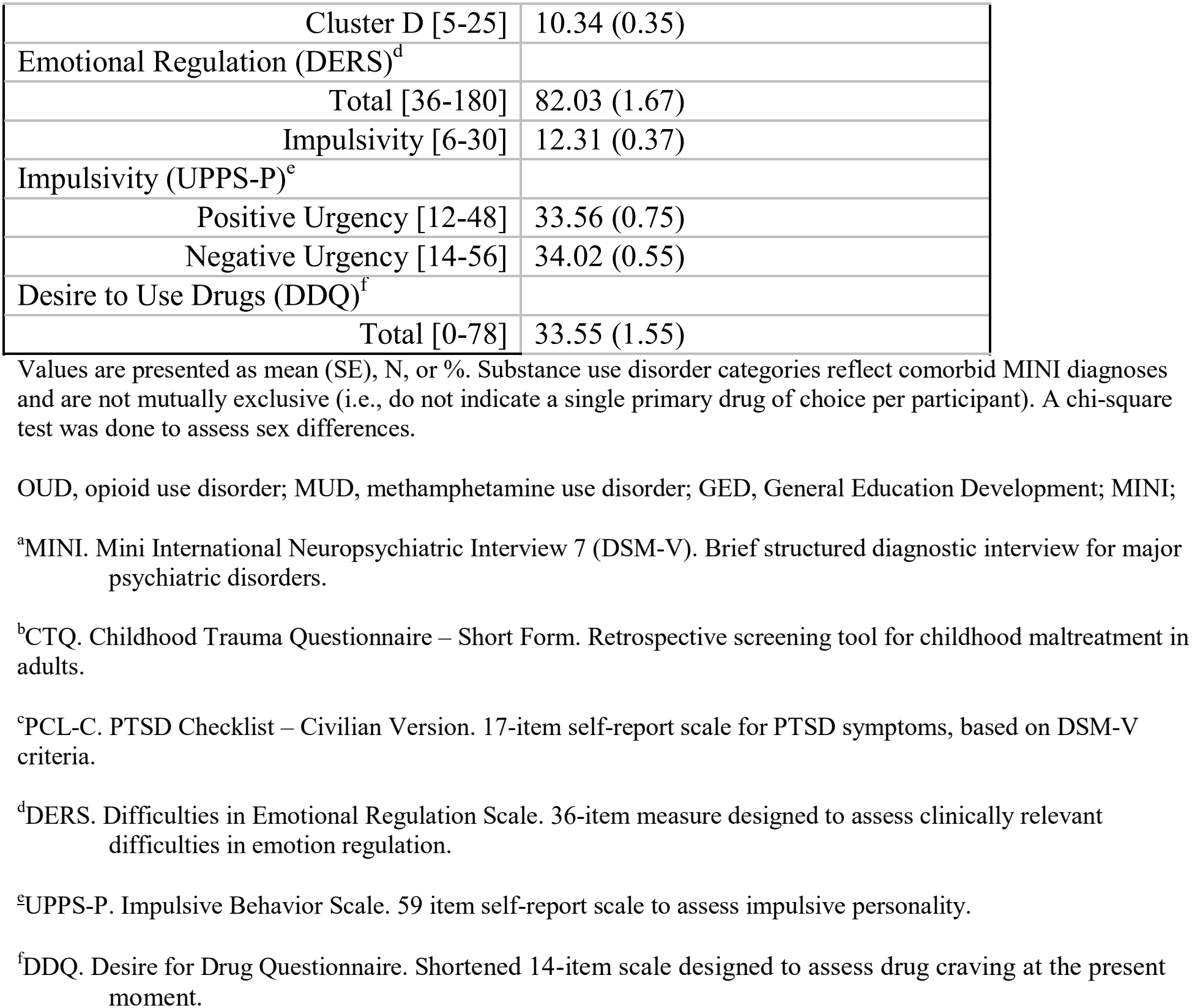
Demographic and clinical characteristics of the study sample.

### Measures

Questionnaires of primary interest for the present analyses included assessments of: (1) traumatic history using the Childhood Trauma Questionnaire – Short Form (CTQ; (37)); (2) PTSD symptomology using the PTSD Checklist-Civilian Version (PCL-C; (38)); (3) emotion regulation using the Difficulties in Emotion Regulation Scale (DERS; (39)); (4) impulsivity using the UPPS-P Impulsive Behavior Scale (UPPS-P; (40)); and (5) drug craving using the Desire for Drug Questionnaire (DDQ; (41)).

### MID Task and Neuroimaging Acquisition

Participants completed two runs (562s, 45 trials each; 90 trials total) of the MID task (6) during fMRI scanning (**Figure S1A**). Each trial began with a fixation period (2–6s), followed by a 2-second anticipation cue signaling the potential to gain (circle) or lose (square) money across six trial types (**Figure S1B**): No Loss ($0.00), Small Loss (−$1.00), Large Loss (−$5.00), No Gain (+$0.00), Small Gain (+$1.00), and Large Gain (+$5.00). The degree of potential gain or loss was indicated by the position of a horizontal line within the cue shape and accompanying text. A target stimulus requiring a rapid button press appeared after a delay of 2.25–3s, followed by a post-response blank (4s) and 2-second feedback indicating the monetary outcome. Task difficulty was calibrated based on practice reaction times (RT) collected prior to scanning, such that participants succeeded on approximately two-thirds of trials, and participants were paid their earnings (∼$30) post-scan. This design allowed dissociation between anticipatory (2-second anticipation cue presentation) and consummatory (2-second feedback presentation) phases of reward and loss processing; the present analyses focused on the anticipatory phase (see Statistical Analyses).

Images were acquired using a GE MRI750 3T scanner. Functional images consisted of 281 contiguous echo-planar imaging (EPI) volumes [TR/TE = 2000/27 ms, FOV/slice = 240/2.9 mm, 128 × 128 matrix, 39 axial slices]. High-resolution structural images were acquired through a 3D axial T1-weighted magnetization-prepared rapid acquisition with gradient-echo sequence [TR/TE = 5/2.012 ms, FOV/slice = 240 × 192/0.9mm, 186 axial slices]. Mean RT per condition was extracted for further group analyses.

### Neuroimaging Data Processing

Neuroimaging data analyses were performed with Analysis of Functional NeuroImaging (AFNI, http://afni.nimh.nih.gov) software (42). The first 3 EPI volumes were discarded for signal equilibrium and scanning noise adaptation. Data were then despiked, slice-timing corrected, co-registered to anatomical volumes, motion corrected, smoothed (4 mm^3^ full width at half maximum), and normalized to standard Montreal Neurological Institute space (resampling voxel size = 2 × 2 × 2 mm^3^).

### Regions of Interest (ROIs)

Regions of interest were extracted using the Brainnetome Atlas (43). Neuroimaging analyses were completed both unilaterally and bilaterally, with bilateral ROIs constructed by averaging left and right hemisphere values. Due to the absence of difference between hemispheres, all analyses included were performed on the following ROIs (**Table S2**): (1) mPFC (both dorsal and ventral); (2) dlPFC; (3) NAc; (4) caudate; (5) putamen; (6) anterior insula; (7) ACC; (8) PCC; and (9) amygdala.

### Statistical Approach

All analyses were conducted in MATLAB version R2024a and 2025b.

#### Behavior

To examine whether trauma-related measures were associated with task engagement, mean RTs were computed for each of the six MID cue conditions (no, small, and large gains and losses) per participant.

#### Single Subject fMRI Analyses

A general linear model (GLM) was employed to analyze single-subject data. Boxcar regressors defined for each participant modeled the blood-oxygen-level dependent (BOLD) response to the anticipation window (4s) in six conditions (15 trials per condition per run): no, small, and large gains and losses. Percent BOLD signal change from baseline was also extracted for: (1) large gains; (2) large losses; and (3) the average of no gains and no losses. Neural responses were modeled during the 2-second cue presentation period (i.e., from cue onset to target appearance), reflecting the anticipatory phase of each trial; inter-trial fixation periods served as the implicit baseline (44). All task regressors were convolved with a canonical hemodynamic response function. In addition, each GLM included nuisance regressors estimated during preprocessing: the first four polynomial terms and six-motion-correction parameters.

#### Group fMRI Analyses

Anticipatory percent BOLD signal change was averaged across all six MID cue conditions for each ROI, yielding a single mean activation value per participant per region. Pearson correlations were then computed between these mean BOLD values and CTQ total scores and PCL-C total scores, respectively, consistent with the individual-differences approach used in prior work relating continuous clinical measures to reward-related neural responses (44). Whereas the Pearson correlations characterize the overall association between trait-level clinical measures and mean regional activation, the linear mixed-effects models were used to additionally model the repeated-measures structure of the MID task, allowing anticipatory activation to be examined across the full range of incentive levels within each ROI rather than relying on a single averaged value. For each ROI, we fit separate linear mixed-effects models (LMEs) with BOLD signal as the dependent variable and MID incentive level (gain: none, small, large; loss: none, small, large) as the primary fixed effect, with CTQ and PCL-C total scores included as continuous predictors to examine their influence on anticipatory activation across gain and loss conditions. As sensitivity analyses, we additionally tested whether drug-of-choice group (opioid vs. methamphetamine use disorder) and in-scanner head motion accounted for CTQ-BOLD associations (see Results). False discovery rate (FDR) correction was applied separately within each family of comparisons: across cue conditions within each ROI for the LME models, and across the full set of ROIs for each clinical measure in the correlation analyses (Table S3, Figure S4).

## Results

### Demographics and Clinical Characteristics

This gender-balanced (52% female) sample reported variable levels of education, from no college, n=88, 58.7%; some college, n=57, 38%; to a college degree: n=5, 3.3%. All participants reported at least one SUD diagnosis, with the most common being methamphetamine use disorder (53%) and opioid use disorder (33%). Most prevalent comorbid psychiatric disorders were major depressive disorder (39%) and generalized anxiety disorder (23%; see **Table 1**). Our sample reported a wide range of childhood traumatic exposure (CTQ: 25-110), with a median score of 43 indicating exposure to at least low severity childhood trauma, and 71% reporting at low-severity exposure. Using published severity cutoffs for the CTQ short form (37), moderate-to-severe exposure was reported for emotional abuse (35.4%; 21.8% severe), physical abuse (30.6%; 21.1% severe), sexual abuse (26.5%; 17.7% severe), emotional neglect (24.5%; 12.9% severe), and physical neglect (29.3%; 15.0% severe). In addition, current PTSD symptomatology (PCL-C) ranged from 17-70 with a median score of 34 indicating probable PTSD diagnosis and 63% reporting at least probable PTSD (using a PCL-C cutoff of ≥31) (**Figure S2**).

### Clinical Correlations

CTQ and PCL-C scores were not significantly correlated with one another (*r* = 0.11, p = 0.18). Similarly, CTQ scores did not differ significantly between participants meeting criteria for probable self-reported PTSD (PCL-C ≥31) and those who did not (t(144) = 1.07, p = 0.29), indicating that childhood trauma exposure and current self-reported PTSD symptom status were dissociable in this sample. However, higher PCL-C scores were associated with higher scores on: (1) DERS total (*r* = 0.56, *p*_FDR_ < 0.001); (2) UPPS-P Negative Urgency (*r* = 0.41, *p*_FDR_ < 0.001); (3) UPPS-P Positive Urgency (*r* = 0.29, *p*_FDR_ < 0.001); and (4) DDQ Total (*r* = 0.41, *p*_FDR_ < 0.001). In contrast, CTQ scores were not significantly correlated with these measures (all *p*_FDR_ > 0.27; **Figure 1, Table 2**).

**Table 2.** Correlations between trauma measures and clinical outcomes.

| **Measure** | **Variable** | **r** | **p** | **p<sub>FDR</sub>** |
|---|---|---|---|---|
| PCL-C Total | DERS Total | 0.556 | **<0.001** | **<0.001** |
|  | UPPS Negative Urgency | 0.410 | **<0.001** | **<0.001** |
|  | UPPS Positive Urgency | 0.293 | **<0.001** | **<0.001** |
|  | DDQ Total | 0.412 | **<0.001** | **<0.001** |
| CTQ Total | DERS Total | 0.067 | 0.423 | 0.564 |
|  | UPPS Negative Urgency | 0.078 | 0.352 | 0.564 |
|  | UPPS Positive Urgency | 0.046 | 0.581 | 0.564 |
|  | DDQ Total | 0.091 | 0.272 | 0.581 |

**Figure 1.**
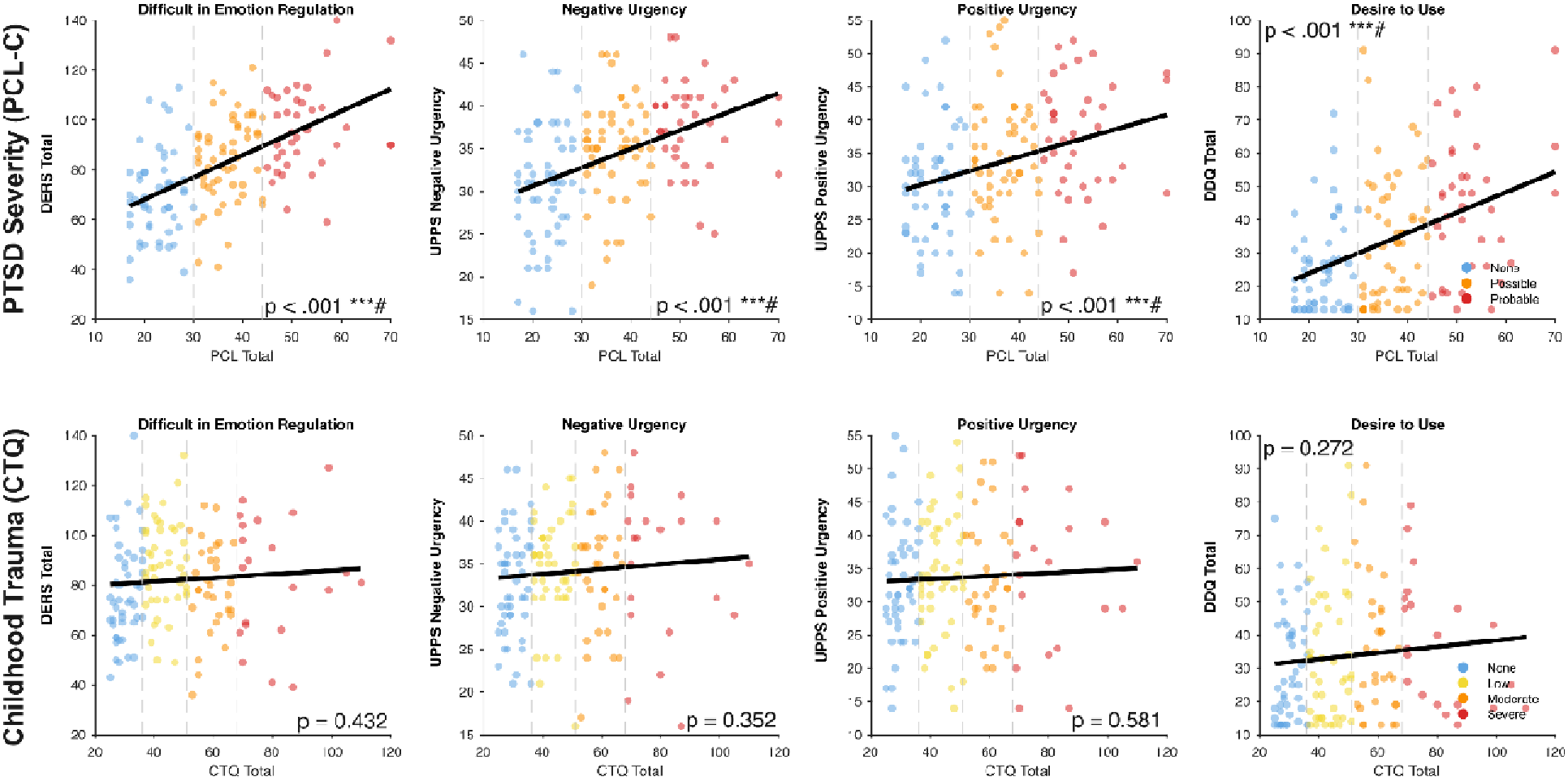
Scatter plots depicting the relationship between PTSD severity (PCL-C Total; top row) and childhood trauma (CTQ total; bottom row) with emotion dysregulation (DERS Total), negative urgency (UPPS Negative Urgency), positive urgency (UPPS Positive Urgency), and desire to use (DDQ Total), stratified by trauma severity (PCL: none, low, moderate, severe; CTQ: none, possible, probable).

We additionally examined general anxiety and depressive symptom severity (PROMIS Anxiety and Depression T-scores) in relation to trauma measures: PCL-C scores were strongly correlated with both PROMIS Anxiety (r = 0.43, p_FDR_ < .001) and PROMIS Depression (r = 0.42, p_FDR_ < .001), whereas CTQ scores showed no significant association with either measure (both p_FDR_ = .41), consistent with trauma-exposure severity and current symptom burden reflecting partially distinct constructs. A direct measure of chronicity of substance use was not available in this dataset; however, a substance-use severity measure was available and was examined as a covariate of the primary CTQ-BOLD associations (see Discussion).

### MID Task

Behaviorally, participants showed the expected pattern of faster RTs for larger monetary cues regardless of valence (**Figure S1C**). Amongst ROIs, all regions exhibited the typical U-shaped BOLD response curve expected from the MID task (Knutson et al., 2001), with greater activation for large gain and loss conditions relative to no-incentive trials (see **Figure S3**). Neither trauma nor clinical measures had any influence on RT.

PCL-C scores were not associated with BOLD signal in any ROI (all *p*_FDR_ > 0.05; see **Table S3, Figure S4**). In contrast, CTQ total scores were negatively associated with BOLD signal across striatal and limbic regions during gain and loss anticipation, with greater CTQ scores predicting blunted responses in the NAc (*r* = -0.21, *p*_FDR_ = 0.049), caudate (*r* = -0.20, *p*_FDR_ = 0.049), amygdala (*r* = -0.20, *p*_FDR_ = 0.049), and ACC (*r* = -0.20, *p*_FDR_ = 0.049) **Table S3; Figure S4**). Higher CTQ scores also related to blunted responses in the dmPFC (*r* = -0.17, *p* = 0.04) and anterior insula (*r* = -0.17, *p* = 0.03), but did not survive FDR correction thresholds (**Table S3, Figure S4**). When grouped into severity levels (none, low, moderate, and severe), a pattern emerged in which increasing childhood trauma severity was associated with increasingly lower BOLD responses across all six ROIs (**Figure 2**).

**Figure 2.**
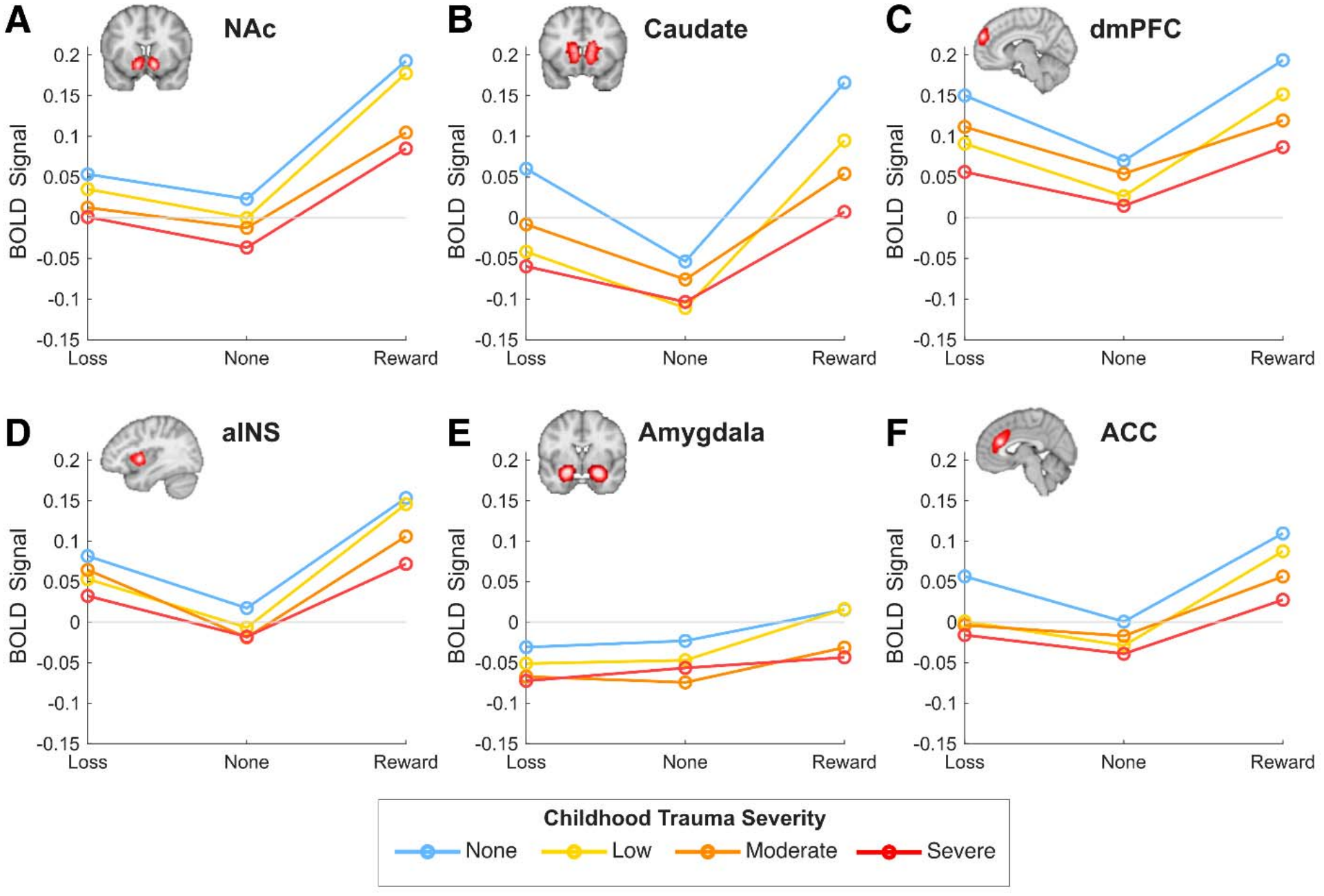
Mean BOLD signal across six trauma-sensitive regions ([**A**] Nucleus accumbens (NAc), [**B**] caudate, [**C**] dorsomedial prefrontal cortex (dmPFC), [**D**] anterior insula (aINS), [**E**] amygdala, and [**F**] anterior cingulate cortex (ACC) as a function of cue valence (Loss, None, Gain) and childhood trauma severity group (CTQ severity score; None, Low, Moderate, Severe). Lines represent group-averaged neural activation. Rates of childhood trauma [**G**] in the study sample. Bottom panel shows Childhood Trauma Questionnaire (CTQ) severity levels. Bars represent the number of participants in each category. Proportions based on sample with scores available. CTQ scores missing for N=3 participant.

To determine whether these associations were independent of general negative affectivity and depressive symptoms, we repeated these correlations controlling for BFI Neuroticism and PHQ-9 depression severity in the subset of participants with complete covariate data (n = 137). The negative association between CTQ scores and BOLD signal remained significant after covarying for both neuroticism and depression severity across all six ROIs (NAc: r = -0.23, pFDR = 0.021; caudate: r = -0.20, pFDR = 0.021; amygdala: r = -0.23, pFDR = 0.021; ACC: r = -0.20, pFDR = 0.023; dmPFC: r = -0.21, pFDR = 0.021; anterior insula: r = -0.21, pFDR = 0.021), indicating that the blunting associated with childhood trauma exposure could not be accounted for by co-occurring depressive symptoms or trait negative affectivity alone.

Because this sample included individuals with both opioid and methamphetamine use disorders, we additionally tested whether drug-of-choice group (opioid, n = 90; methamphetamine, n = 57) accounted for the association between CTQ scores and BOLD signal. Covarying for group did not meaningfully change the strength of the association for any ROI (partial r range: -0.18 to -0.22, all p < .04). Examined separately by group, correlations were numerically stronger among participants with opioid use disorder (r range: -0.20 to -0.28) than among those with methamphetamine use disorder (r range: -0.09 to -0.18, not significant at this smaller subsample size), though the direction and approximate magnitude of the association were consistent across both groups, suggesting the overall finding was not driven by a single drug-of-choice subgroup.

We also tested whether in-scanner head motion accounted for these associations. Controlling for mean framewise motion during the MID task did not attenuate the relationship between CTQ scores and BOLD signal in any ROI; associations remained significant after FDR correction across all six ROIs (NAc: r = -0.22, pFDR = 0.020; caudate: r = -0.21, pFDR = 0.020; amygdala: r = -0.23, pFDR = 0.020; ACC: r = -0.20, pFDR = 0.020; dmPFC: r = -0.18, pFDR = 0.027; anterior insula: r = -0.19, pFDR = 0.023), indicating that the blunting associated with childhood trauma exposure was not attributable to differences in head motion.

## Discussion

The present study examined neural responses to monetary gain and loss anticipation in treatment-enrolled individuals with SUD, primarily methamphetamine and/or opioid use disorders, using the MID task during fMRI, to identify how childhood trauma severity and current PTSD symptom burden relate to anticipatory reward circuit function. This work addresses two critical gaps in the literature: (1) the underrepresentation of methamphetamine and opioid use disorders in MID neuroimaging research; and (2) the largely untested hypothesis that trauma and SUD share overlapping patterns of blunted frontostriatal reward anticipation.

We did not include a non-SUD comparison group in this study, and therefore could not examine whether MID task activation differed between individuals with SUD and healthy controls; this sample was designed to characterize heterogeneity in trauma- and PTSD-related reward processing within SUD rather than to establish SUD-related differences from a comparison group. We did, however, examine whether severity of substance use accounted for the observed associations. Using a substance-use severity measure (PROMIS Substance Use Severity T-score; M = 55.3, SD = 7.0), the association between CTQ scores and BOLD signal remained significant after covarying for severity of use across all six ROIs (NAc: r = -0.20, pFDR = 0.026; caudate: r = -0.20, pFDR = 0.026; amygdala: r = -0.21, pFDR = 0.026; ACC: r = -0.20, pFDR = 0.026; dmPFC: r = -0.17, pFDR = 0.038; anterior insula: r = -0.19, pFDR = 0.031), indicating that the blunting associated with childhood trauma exposure was not attributable to differences in substance use severity.

Our central findings revealed that: (1) childhood trauma and current self-reported PTSD symptoms did not share significant variance across individuals with SUD; (2) greater current PTSD symptom severity (but not childhood trauma) was associated with greater emotion dysregulation, emotional impulsivity, and current drug craving; and (3) greater childhood trauma exposure (but not current PTSD symptom severity) was associated with lower striatal, amygdalar, and ACC BOLD signal to cues signaling probable monetary attainment or avoidance of loss.

This suggests that PTSD symptom burden shapes the affective and motivational aspects of daily functioning, increasing reactivity and craving, without necessarily disrupting the neural mechanisms engaged during structured incentive processing. Importantly, the absence of observed activation differences associated with current PTSD symptoms during reward anticipation should not be interpreted as evidence that PTSD is neurobiologically inconsequential. Prior work has demonstrated PTSD-related alterations in resting-state connectivity, gray matter volume, and default mode network functions ((45); (46)), suggesting that symptom-related neural differences may manifest across multiple levels of brain organization beyond task-evoked activation. Alternatively, current PTSD symptoms may influence functioning during processes not well captured by this task, such as the regulation of explicitly emotional information. The influence of current PTSD symptoms on substance use behavior may operate primarily through affective and motivational channels that are most apparent in naturalistic drug-cue environments rather than controlled laboratory tasks. Additionally, we lack data on symptom duration and cannot determine whether longer-standing symptoms would be more likely to produce circuit-level reorganization detectable via activation-based fMRI measures of reward anticipation.

The prominence of early-life adversity effects aligns with evidence that it produces lasting reorganization of corticostriatal and limbic circuits involved in affective valuation and reinforcement learning (32), alterations that may predate substance use onset and contribute to long-term vulnerability. Early-life adversity occurs during sensitive periods of heightened neuroplasticity, which may explain why its corticostriatal effects are more deeply embedded and developmentally cumulative than those associated with current PTSD burden (47). Notably, the finding that caudate activation was influenced by childhood trauma exposure suggests that early life adversity may shift reinforcement learning toward rigid, habitual patterns less sensitive to outcome value (48), a profile that may make it harder to disengage from substance use when consequences become negative, increasing risk for escalation and relapse.

This double dissociation, clinical burden without neural disruption during reward/loss anticipation in current PTSD symptomology, and neural disruption without clear evidence of clinical burden, at least on the measures assessed here, in childhood trauma, underscores the importance of distinguishing the timing and type of trauma exposure (i.e., early-life trauma history versus current posttraumatic symptomatology) when characterizing reward dysfunction in SUD. It also highlights that trauma screening should be a standard component of neuroimaging research in SUD populations, as failure to account for trauma history may obscure meaningful sources of neural variability and limit the translational value of reward circuit findings. Together, these findings support a model in which trauma typology produces distinct profiles of reward circuit dysfunction through different mechanisms and timescales, both carrying direct implications for trauma-informed treatment but likely requiring distinct intervention targets.

Notably, childhood trauma severity was associated with blunted anticipatory BOLD signal to both gain and loss cues, rather than selectively to one valence. This bidirectional blunting suggests a more general dampening of incentive salience, rather than a valence-specific deficit in approach or avoidance motivation, and may reflect a broader reduction in the capacity of monetary incentives to engage these circuits regardless of whether they signal possible reward or possible loss. Clinically, this pattern raises the possibility that individuals with greater childhood trauma exposure may be less responsive to incentive-based behavioral interventions generally, rather than selectively under-responsive to reward, suggesting that treatments relying on reinforcement or loss-avoidance framing alone may need to be paired with approaches that more directly target salience and motivation.

Although Generalized Anxiety Disorder was diagnosed more frequently than PTSD in this sample (23.3% vs. 4.7% by MINI diagnosis), we focused on childhood trauma and PTSD symptomatology rather than general anxiety given the specificity of the proposed mechanistic pathway: childhood adversity and PTSD are directly implicated in stress-sensitization of mesolimbic and corticostriatal circuitry, producing a plausible and well-characterized route to reward-circuit dysfunction, whereas comparable reward-specific mechanisms for generalized anxiety are less established. Notably, the relatively low rate of categorical PTSD diagnosis in this sample stood in contrast to a substantially higher rate of clinically meaningful self-reported PTSD symptoms (63% meeting probable PTSD thresholds on the PCL-C), suggesting that symptom-level trauma burden, rather than categorical diagnosis alone, captures population-relevant variability that would otherwise be overlooked. Moreover, because childhood trauma frequently precedes the later development of anxiety and stress-related symptomatology, examining trauma directly offers a developmentally upstream vantage point on reward circuit disruption in SUD, one that may partially underlie elevated rates of several comorbid anxiety presentations rather than operating independently of them.

Several limitations warrant consideration. The cross-sectional design precludes causal inference about the directionality of associations between trauma and reward circuit function. Additionally, the CTQ captures retrospective self-report of childhood experiences and is subject to recall biases (49). Future work using prospective or longitudinal designs would strengthen inference about developmental effects of early trauma on outcome processing in SUD populations.

## Acknowledgements and Disclosures

MCMG acknowledges support from NIGMS (Grant number: P20GM121312; PI: M.P. Paulus). This project was a secondary analysis of data from a study conceived by Martin P. Paulus and financially supported by the Laureate Institute for Brain Research (Grant number: LIBR # 2018-012-00). The findings and conclusions of this article are those of authors and do not necessarily represent the views and opinions of these organizations, which had no role in designing, or planning the study, in collecting, analyzing, or interpreting the data, in writing the article, or deciding to submit it for publication.

The fMRI findings with childhood trauma were presented by MCMG in poster format at Society for Biological Psychiatry conference in New York City, NY, 2026.

SAA and SIQ accessed the data, performed data cleaning, helped create the analysis pipeline, performed data analysis, contributed to result interpretation. They equally contributed to the manuscript by providing the first draft of the article and editing subsequent drafts. RK performed fMRI data preprocessing and assisted with first and second-level analyses. JLS and RLA edited and reviewed subsequent drafts of the manuscript. MCMG guided and oversaw data analysis, helped create and reviewed the analysis pipeline, helped with result interpretation, edited drafts of the articles, and provided supervision and guidance. All authors made substantive contributions to the article, read, and approved the final article.

The authors report no biomedical financial interest, or potential conflicts of interest.

