## Supplemental for "Childhood Trauma Severity Compounds Reward Anticipation Disruption in Substance Use Disorders"

**Table S1: Demographics and Comorbid Clinical Diagnoses**

| Sample Description | Total Sample (N = 150) |
| --- | --- |
| DEMOGRAPHICS | |
| Race |  |
| American Indian | 10 (6.7%) |
| Black | 6 (4%) |
| Multiracial | 20 (13.3%) |
| White | 114 (76%) |
| Ethnicity |  |
| Hispanic/Latino | 6 (4%) |
| Not Hispanic/Latino | 144 (96%) |
| Education |  |
| < High School | 34 (22.7%) |
| High School Graduate/GED | 54 (36%) |
| Some College/Tech School | 57 (38%) |
| Bachelors | 4 (2.7%) |
| Graduate Degree | 1 (0.7%) |
| Employment |  |
| Employed | 25 (16.7%) |
| Unemployed | 115 (76.7%) |
| Unknown | 10 (6.7%) |
| PSYCHIATRIC DISORDERS | |
| MOOD DISORDERS |  |
| Bipolar Disorder | 10 (6.7%) |
| Bipolar Type I | 8 |
| Bipolar Type II | 2 |
| Major Depressive Disorder | 59 (39.3%) |
| ANXIETY DISORDERS |  |
| Agoraphobia | 17 (11.3%) |
| Obsessive Compulsive Disorder | 7 (4.7%) |
| Panic Disorder | 7 (4.7%) |
| Social Phobia | 25 (16.7%) |
| Social Anxiety Disorder | 2 (1.3%) |
| Binge Eating Disorder | 10 (6.7%) |
| SUBSTANCE USE DISORDERS |  |
| Amphetamine Use Disorder | 79 (52.6%) |
| Mild | 1 |
| Moderate | 4 |
| Severe | 72 |
| Dependence | 2 |
| Opioid Use Disorder | 49 (32.7%) |
| Mild | 1 |
| Moderate | 1 |
| Severe | 43 |
| Dependence | 4 |
| Alcohol Use Disorder | 21 (14%) |
| Mild | 4 |
| Moderate | 2 |
| Severe | 12 |
| Dependence | 3 |
| Cocaine Use Disorder | 4 (2.7%) |
| Mild | 0 |
| Moderate | 1 |
| Severe | 3 |
| Dependence | 0 |
| Cannabis Use Disorder | 43 (28.7%) |
| Mild | 11 |
| Moderate | 12 |
| Severe | 19 |
| Dependence | 1 |
| Hallucinogen Use Disorder | 3 (2%) |
| Mild | 1 |
| Moderate | 0 |
| Severe | 2 |
| Sedative Use Disorder | 17 (11.3%) |
| Mild | 1 |
| Moderate | 2 |
| Severe | 12 |
| Dependence | 2 |

Values are presented as N (%) or Mean (SD). Diagnoses assessed via MINI International Neuropsychiatric Interview 7 (DSM-5).

**Table S2. Bilateral Regions of Interest Defined Using the Brainnetome Atlas**

| **Region of Interest** | **Abbreviation** | **Brainnetome Parcels** |
| --- | --- | --- |
| Nucleus Accumbens | NAc | 223, 224 |
| Ventromedial Prefrontal Cortex | vmPFC | 41, 42, 45, 46, 47, 48, 49, 50, 187, 188 |
| Dorsomedial Prefrontal Cortex | dmPFC | 1, 2, 11, 12 |
| Dorsolateral Prefrontal Cortex | dlPFC | 3, 4, 15, 16, 17, 18, 23, 24, 25, 26 |
| Anterior Insula | aINS | 165, 166, 167, 168 |
| Anterior Cingulate Cortex | ACC | 177, 178, 179, 180, 183, 184 |
| Amygdala | — | 211, 212, 213, 214 |
| Caudate | Caudate | 219, 220, 227, 228 |
| Putamen | Putamen | 225, 226, 229, 230 |
| Posterior Cingulate Cortex | PCC | 153, 154, 175, 176, 181, 182 |

***Table S2.*** A priori regions of interest selected from preexisting literature. Hemispheres were analyzed bilaterally.

**Table S3. Combined MID cue BOLD correlation with trauma measures.**

|  | **Posttraumatic Symptoms (PCL)** | | | **Childhood Traumatic Exposure (CTQ)** | | |
| --- | --- | --- | --- | --- | --- | --- |
| **ROI** | **r-value** | **p-value** | **q-value** | **r-value** | **p-value** | **q-value** |
| NAc | 0.027 | 0.74 | 0.92 | -0.21 | **0.011** | **0.049** |
| Caudate | 0.0028 | 0.97 | 0.98 | -0.20 | **0.013** | **0.049** |
| Putamen | -0.035 | 0.67 | 0.92 | -0.15 | 0.071 | 0.10 |
| vmPFC | -0.025 | 0.76 | 0.92 | -0.101 | 0.20 | 0.24 |
| dmPFC | -0.037 | 0.652 | 0.92 | -0.17 | **0.036** | 0.072 |
| dlPFC | -0.091 | 0.27 | 0.92 | -0.15 | 0.068 | 0.10 |
| aINS | -0.047 | 0.57 | 0.92 | -0.17 | **0.033** | 0.072 |
| Amygdala | -0.18 | **0.029** | 0.34 | -0.20 | **0.016** | **0.049** |
| ACC | -0.0018 | 0.98 | 0.98 | -0.20 | **0.016** | **0.049** |
| PCC | -0.10 | 0.21 | 0.92 | -0.14 | 0.079 | 0.10 |

***Table S3.*** Bold values indicate significance. Correlations between trauma measure (PCL-C = PTSD Checklist – Civilian Version; CTQ = Childhood Trauma Questionnaire) and region of interest (ROI).

Because bilateral ROIs were constructed by averaging left and right hemisphere values, we additionally examined these correlations separately by hemisphere to confirm that the bilateral pattern was not driven by one hemisphere. Results were highly similar in direction and magnitude across hemispheres (Tables S4-S5), supporting the use of bilateral ROIs in the main analyses.

**Table S4. Left hemisphere MID cue BOLD correlation with trauma measures.**

|  | **Posttraumatic Symptoms (PCL)** | | | **Childhood Traumatic Exposure (CTQ)** | | |
| --- | --- | --- | --- | --- | --- | --- |
| **ROI** | **r-value** | **p-value** | **q-value** | **r-value** | **p-value** | **q-value** |
| NAc | -0.004 | 0.96 | 0.96 | -0.185 | 0.03 | 0.07 |
| Caudate | 0.017 | 0.84 | 0.96 | -0.185 | 0.03 | 0.07 |
| Putamen | -0.022 | 0.79 | 0.96 | -0.179 | 0.03 | 0.07 |
| vmPFC | -0.019 | 0.82 | 0.96 | -0.110 | 0.18 | 0.20 |
| dmPFC | -0.019 | 0.82 | 0.96 | -0.062 | 0.45 | 0.45 |
| dlPFC | -0.102 | 0.22 | 0.74 | -0.118 | 0.15 | 0.19 |
| aINS | -0.030 | 0.72 | 0.96 | -0.171 | 0.04 | 0.08 |
| ACC | 0.012 | 0.88 | 0.96 | -0.159 | 0.05 | 0.09 |
| Amygdala | -0.186 | 0.03 | 0.25 | -0.231 | **0.005** | **0.05** |
| PCC | -0.115 | 0.17 | 0.74 | -0.131 | 0.11 | 0.16 |

***Table S4.*** Bold values indicate significance. Correlations between trauma measure (PCL-C = PTSD Checklist – Civilian Version; CTQ = Childhood Trauma Questionnaire) and region of interest (ROI), computed separately for the left and right hemisphere parcels comprising each bilateral ROI (n = 147).

**Table S5. Right hemisphere MID cue BOLD correlation with trauma measures.**

|  | **Posttraumatic Symptoms (PCL)** | | | **Childhood Traumatic Exposure (CTQ)** | | |
| --- | --- | --- | --- | --- | --- | --- |
| **ROI** | **r-value** | **p-value** | **q-value** | **r-value** | **p-value** | **q-value** |
| NAc | 0.037 | 0.66 | 0.88 | -0.229 | **0.005** | **0.04** |
| Caudate | -0.029 | 0.73 | 0.88 | -0.220 | **0.007** | **0.04** |
| Putamen | -0.063 | 0.45 | 0.88 | -0.132 | 0.11 | 0.11 |
| vmPFC | -0.016 | 0.85 | 0.88 | -0.134 | 0.11 | 0.11 |
| dmPFC | -0.031 | 0.71 | 0.88 | -0.139 | 0.09 | 0.11 |
| dlPFC | -0.114 | 0.17 | 0.57 | -0.159 | 0.06 | 0.08 |
| aINS | -0.071 | 0.39 | 0.88 | -0.188 | 0.02 | 0.05 |
| ACC | -0.013 | 0.88 | 0.88 | -0.175 | 0.03 | 0.06 |
| Amygdala | -0.184 | 0.03 | 0.26 | -0.183 | 0.03 | 0.05 |
| PCC | -0.135 | 0.10 | 0.52 | -0.211 | **0.01** | **0.04** |

***Table S5.*** Bold values indicate significance. Correlations between trauma measure (PCL-C = PTSD Checklist – Civilian Version; CTQ = Childhood Trauma Questionnaire) and region of interest (ROI), computed separately for the left and right hemisphere parcels comprising each bilateral ROI (n = 147).

**Supplemental Results**

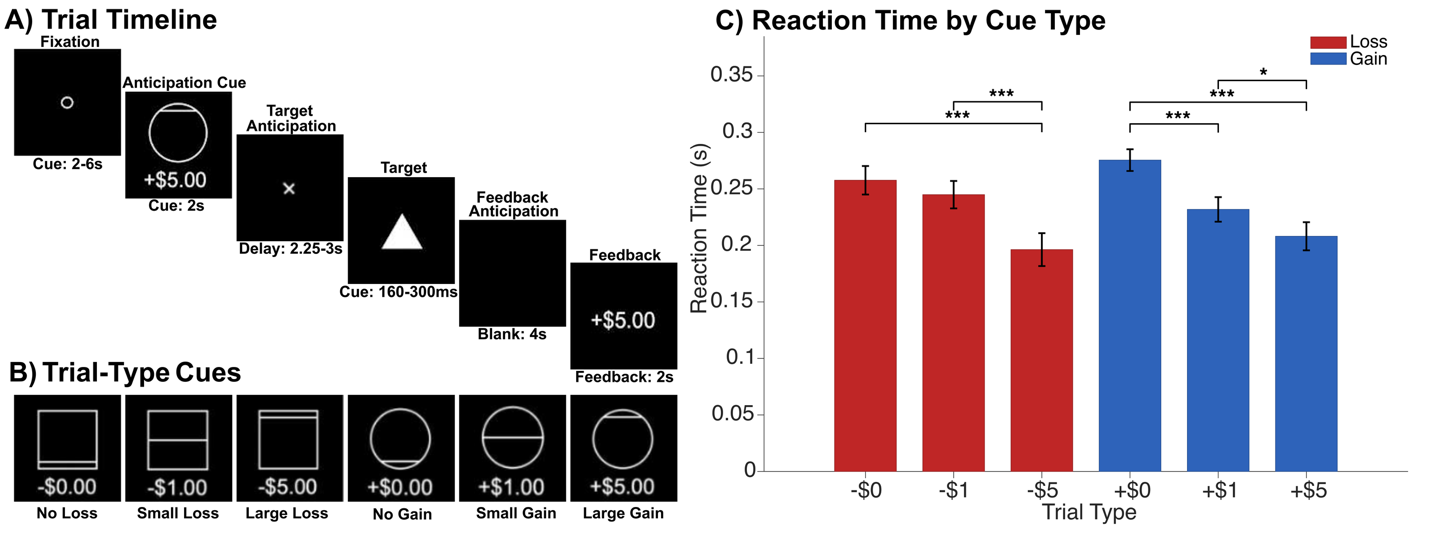
***Figure S1.*** The Monetary Incentive Delay (MID) task. **(A)** Sample trial sequence for a High Gain trial, showing fixation (2–6s), anticipation cue (2s), target anticipation with delay (2.25–3s), post-response blank (ends 4s after target anticipation started), and feedback (2s). **(B)** Squares denote loss trials (No Loss = $0.00, Low Loss = −$1.00, High Loss = −$5.00), and circles denote gain trials (No Gain = +$0.00, Low Gain = +$1.00, High Gain = +$5.00). **(C)** Mean reaction times (seconds) by trial type, demonstrating faster responses for larger monetary outcomes regardless of valence.

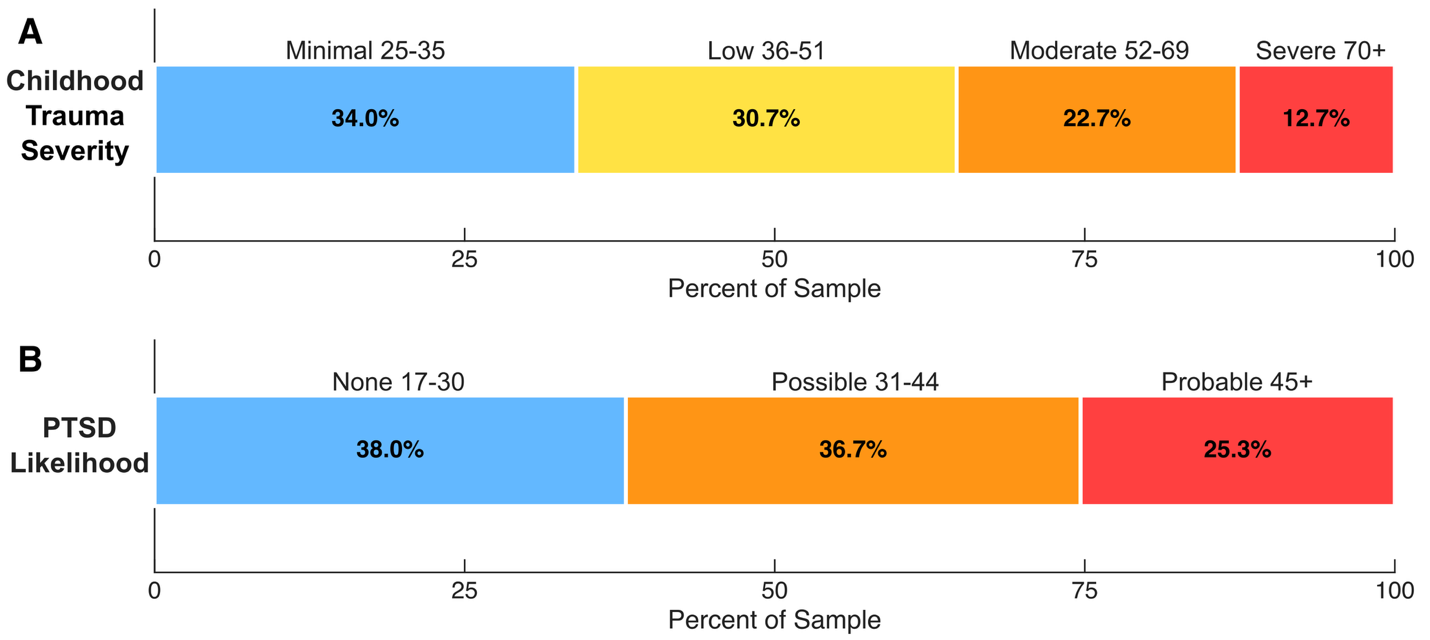
***Figure S2.*** Rates of childhood trauma [**A**] and PTSD symptoms [**B**] in the study sample. Top panel shows Childhood Trauma Questionnaire (CTQ) severity levels; bottom panel shows PTSD symptom severity (PCL-C). Bars represent the number of participants in each category. Proportions based on sample with scores available. CTQ scores missing for N=3 and PCL-C scores missing for N=3 participants.

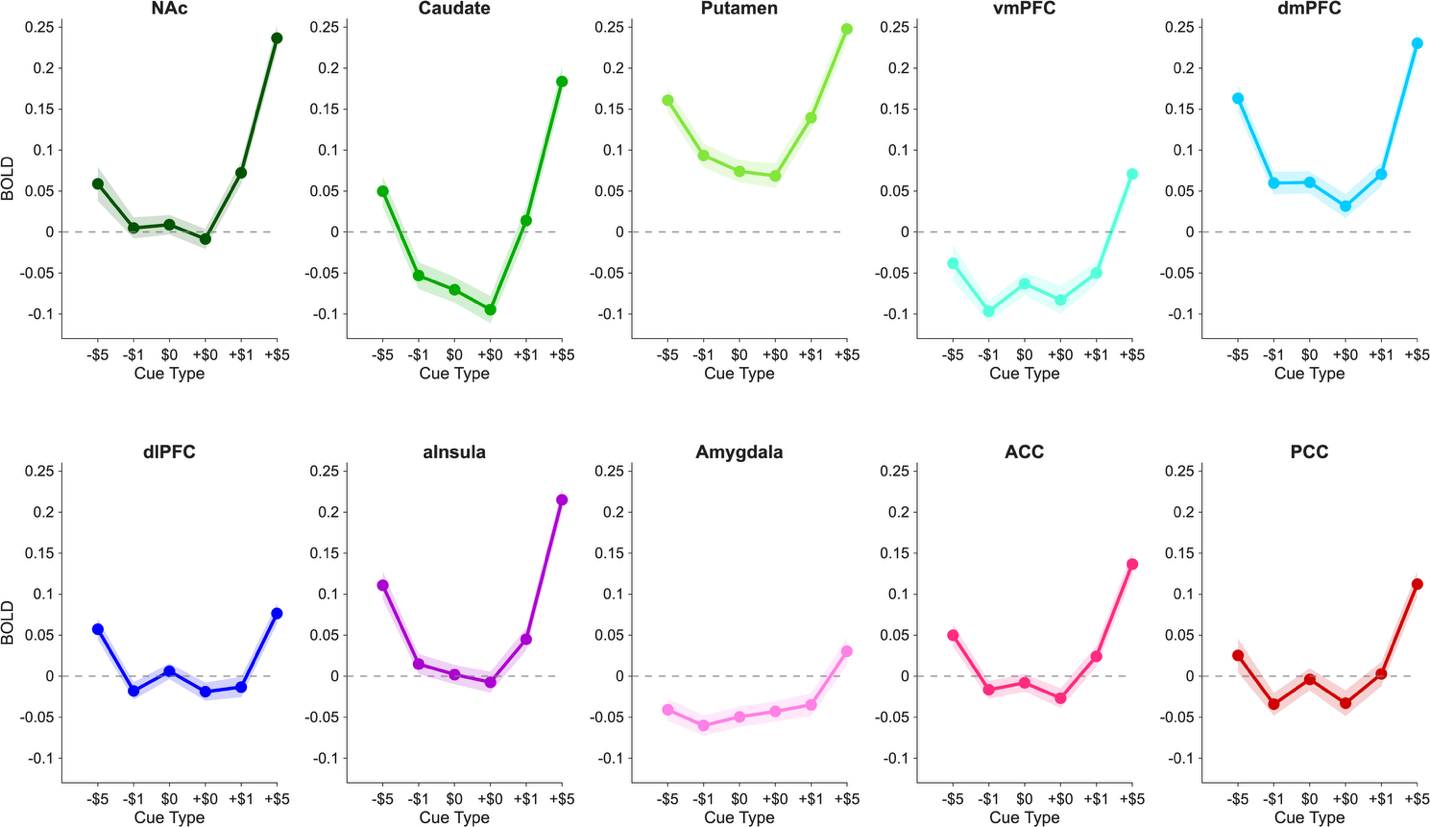
***Figure S3.*** Whole brain analyses were conducted to compare neural activation during gain (-$5, -$1, -$0) and loss ($5, $1, $0) within individuals with substance use disorders (SUDs). U-shaped curves display the BOLD signal across task conditions.

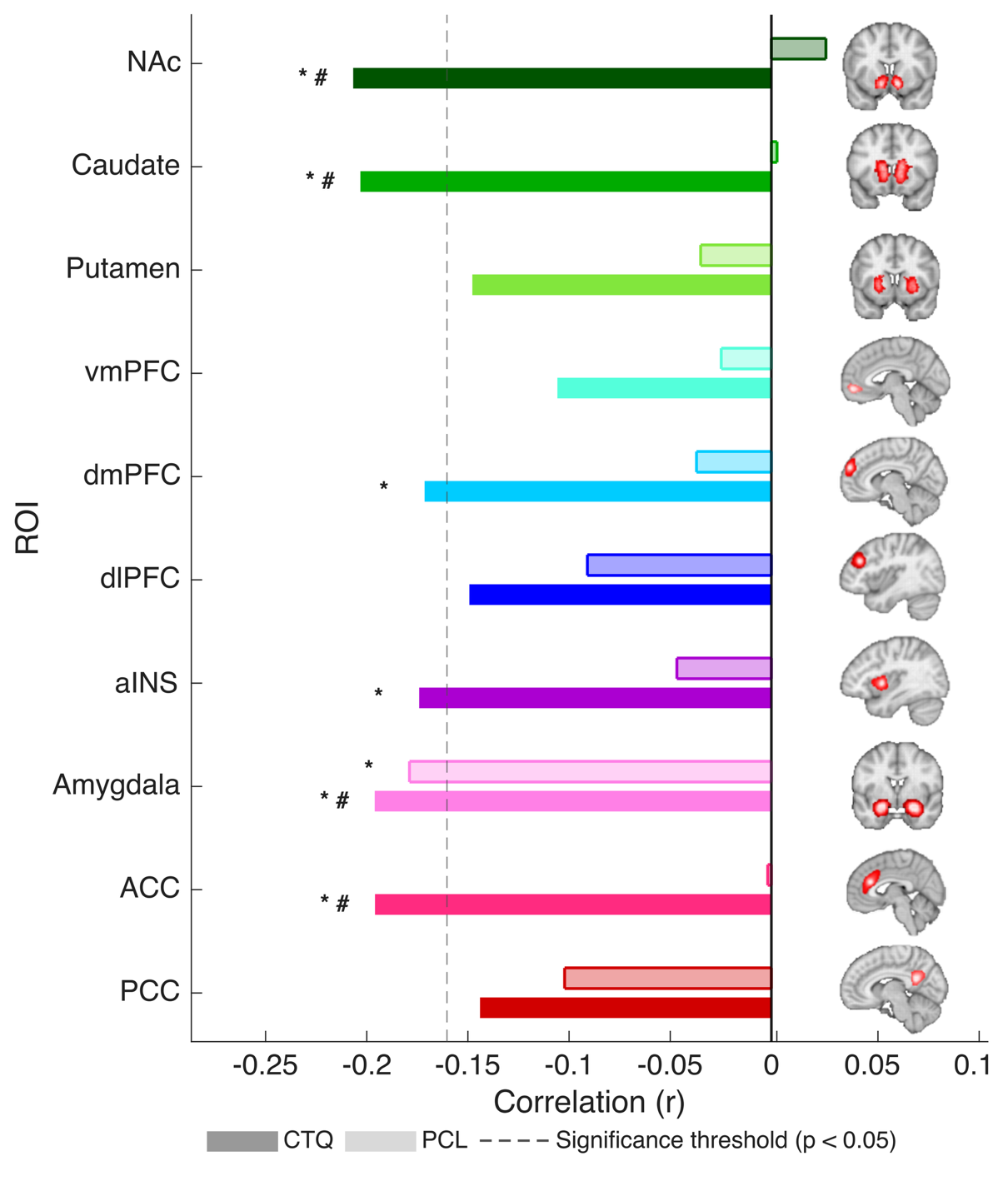

***Figure S4.*** Pearson correlations between mean BOLD activation (averaged across all outcome conditions) and CTQ total scores (lighter tint, dashed border) and PCL-C total scores (solid) for all regions of interest. Negative values indicate that greater trauma exposure or symptom severity was associated with blunted activation. * p < 0.05, ** p < 0.01, *** p < 0.001 (uncorrected). FDR correction is applied for each condition across n=12 ROIs: # p < 0.05 FDR corrected. The dashed line shows the significance threshold for where the correlation is statistically significant (p < 0.05, two-tailed).
